# CENcyclopedia: Dynamic Landscape of Kinetochore Architecture Throughout the Cell Cycle

**DOI:** 10.1101/2024.12.05.627000

**Authors:** Yu-Chia Chen, Ece Kilic, Evelyn Wang, Will Rossman, Aussie Suzuki

**Affiliations:** McArdle Laboratory for Cancer Research, Department of Oncology, University of Wisconsin-Madison, Madison, Wisconsin, USA; Molecular Cellular Pharmacology Graduate Program, University of Wisconsin-Madison, Madison, Wisconsin, USA; Carbone Comprehensive Cancer Center, University of Wisconsin-Madison, Madison, Wisconsin, USA

## Abstract

The kinetochore, an intricate macromolecular protein complex located on chromosomes, plays a pivotal role in orchestrating chromosome segregation. It functions as a versatile platform for microtubule assembly, diligently monitors microtubule binding fidelity, and acts as a force coupler. Comprising over 100 distinct proteins, many of which exist in multiple copies, the kinetochore’s composition dynamically changes throughout the cell cycle, responding to specific timing and conditions. This dynamicity is important for establishing functional kinetochores, yet the regulatory mechanisms of these dynamics have largely remained elusive. In this study, we employed advanced quantitative immunofluorescence techniques to meticulously chart the dynamics of kinetochore protein levels across the cell cycle. These findings offer a comprehensive view of the dynamic landscape of kinetochore architecture, shedding light on the detailed mechanisms of microtubule interaction and the nuanced characteristics of kinetochore proteins. This study significantly advances our understanding of the molecular coordination underlying chromosome segregation.

## Introduction

Chromosome segregation ensures the equal distribution of replicated genomes into daughter cells. Errors in this process can lead to aneuploidy, an abnormal number of chromosomes and a hallmark of cancer. The kinetochore is a multiprotein complex that assembles on the centromeric chromatin and orchestrates chromosome segregation. Kinetochores not only serve as a structural platform for microtubule assembly but also ensure faithful chromosome segregation by actively monitoring kinetochore-microtubule interactions^1^.

The kinetochore architecture can be divided into three regions: inner kinetochore, outer kinetochore, and corona^2–4^. Each region has a unique composition of proteins. The inner kinetochore consists of constitutive centromere-associated network (CCAN) proteins, 16 different subunits that directly assemble on the centromeric chromatin and serve as the foundation of kinetochore assembly^5,6^. The outer kinetochore contains the highly conserved KMN network, which includes the Knl1 complex (Knl1C), the Mis12 complex (Mis12C), and the Ndc80 complex (Ndc80C)^7–9^. Ndc80C is the primary microtubule-binding site at kinetochores, while Knl1C serves as a major platform for the assembly of spindle assembly checkpoint (SAC) proteins to monitor attachment errors. The corona is the outermost layer of kinetochores, where SAC-related and microtubule-associated proteins reside^10^. The primary function of corona proteins is to form a higher-order assembly that facilitates the capture of chromosomes by spindle microtubules^11^.

Recent efforts have led to the identification of over 100 different kinetochore-related proteins, which are dynamically regulated throughout the cell cycle to form functional kinetochores^12^. While some proteins are constitutively assembled at centromeres, others are recruited to kinetochores at specific cell cycle stages^13^. Previous studies on kinetochore protein dynamics have been often focused on a small subset of proteins or their turnover at kinetochores^14–18^, failing to fully capture their abundance and regulation of their recruitment across the entire cell cycle. This limitation hinders the comprehensive understanding of the kinetochore’s dynamic landscape. In this study, we utilized quantitative immunofluorescence (qIF) microscopy to profile the dynamics of 31 kinetochore proteins and 5 mitotic kinase substrates in asynchronous non-transformed RPE1 cells. Our approach provides a holistic view of the kinetochore architecture across all cell cycle phases, including the less-explored G1, S, and G2 phases. We reveal that CCAN proteins are promptly recruited to kinetochores during S phase when centromeres are replicated. Unexpectedly, we observe that Mis12C localizes to kinetochores as early as in the middle/late G1 phase through CENP-C in an Aurora-B-independent manner, while the recruitment of Knl1C and Ndc80C initiates in late S phase. Additionally, the assembly of SAC-related proteins, motor proteins, and kinases at kinetochores follows a multi-step process, characterized by a subtle yet critical time lag in their association and dissociation. This comprehensive analysis of kinetochore protein dynamics provides new insights into the structural and functional organization of kinetochores, contributing to a more complete understanding of the mechanisms that ensure faithful chromosome segregation.

## Results

### Strategies for quantifying kinetochore protein abundance across the cell cycle

To quantitatively define kinetochore protein dynamics throughout the cell cycle, we divided the cell cycle into the following stages: G1, S, G2 (late G2), and the sub-stages of M phase. For precise measurement of protein levels at kinetochores in each cell cycle stage, we employed our recently-developed immunofluorescence-based method for cell cycle stage identification^19^ (Fig. 1a). Specifically, G1 phase cells are defined by the absence CENP-F signals in the nucleus. S phase cells display distinct nuclear PCNA puncta, CENP-F signals, and brighter CENP-C signals. G2 phase cells exhibit uniform PCNA nuclear staining like G1, along with elevated nuclear CENP-F signals. The vast majority of sister kinetochores are within diffraction-limited distances during early G2 phase, whereas distinct paired kinetochore signals emerge prominently in late G2 phase. Since most non-CCAN kinetochore proteins are absent from kinetochores between G1 and early G2 phase, in our kinetochore qIF analysis, most data obtained from these stages were combined into one category, with a subset of kinetochore protein measurements distinguishing between stages in interphase (see **Methods** for details). Sub-stages of M phase were identified based on distinct DNA morphologies (Fig. 1a and Extended Data Fig. 1a,b). To prevent artificial dissociation of kinetochore proteins, we utilized either paraformaldehyde (PFA) or methanol fixation without permeabilization prior to or during fixation, for all qIF experiments, except for Bub3 and Ska3 staining (Extended Data Fig. 1c). For these exceptions, pre-extraction fixation was applied. Detailed information about antibodies and their corresponding fixation method is listed in Supplementary Table 1-3. Signal intensities for target proteins at individual kinetochores were measured using a local background correction method that we previously developed (Extended Data Fig. 1d,e)^20,21^. In the case where target proteins were absent from kinetochores, CENP-C foci served as a reference at the corresponding positions. Notably, we achieved a signal-to-noise (S/N) ratio exceeding 10 for most antibodies (27 out of 36) at their peak levels (Extended Data Fig. 1f), enabling highly sensitive and accurate quantifications.

**Fig. 1.**
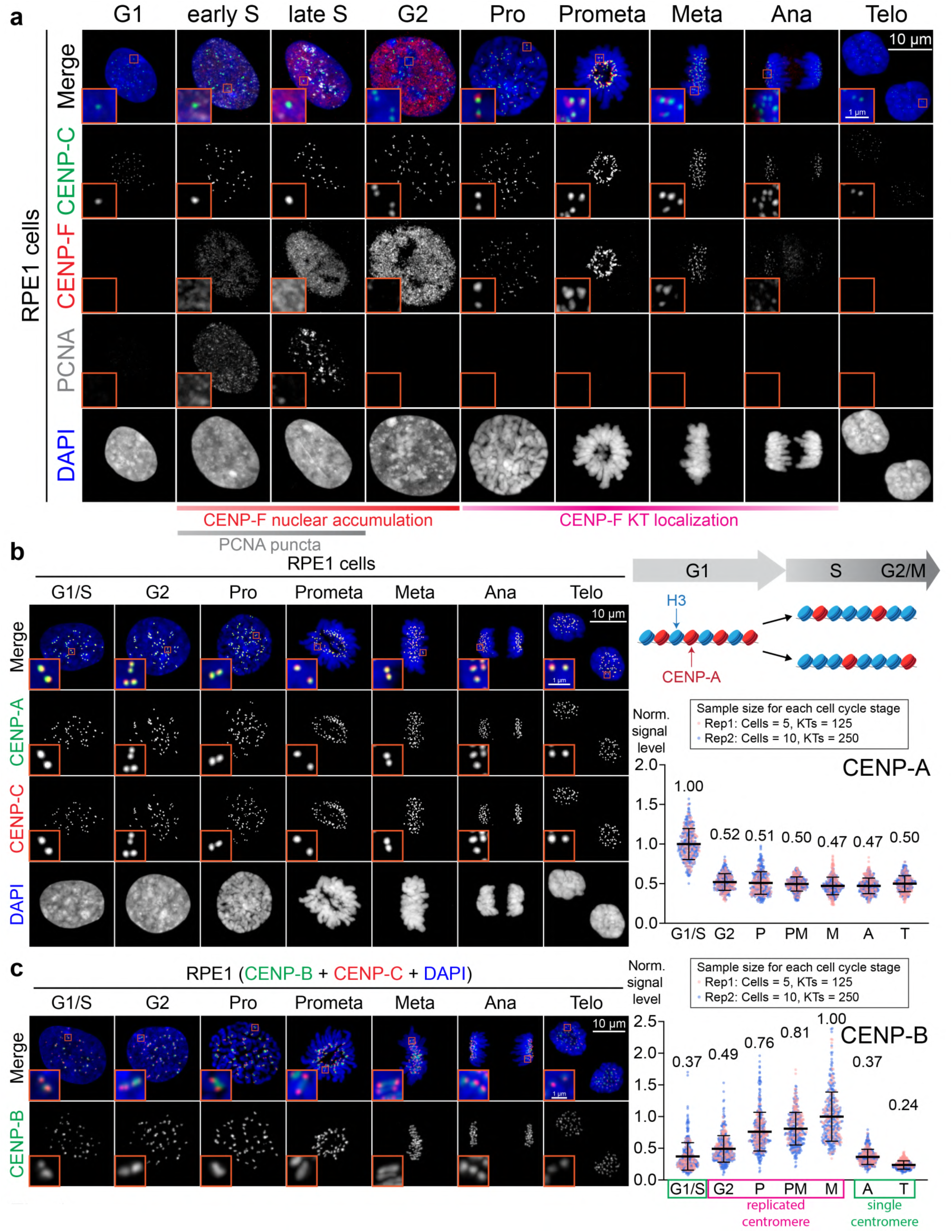
Characterization of the cell cycle stage determination method. **a**, Representative images of CENP-C, CENP-F, PCNA and DNA staining throughout the cell cycle of RPE1 cells. **b**, Left: representative images of CENP-A throughout the cell cycle. CENP-C is used as a kinetochore marker. Top right: schematic diagram of CENP- A distribution at centromeres before and after DNA replication during S phase. Bottom right: quantification of CENP-A levels at kinetochores at different stages of the cell cycle. Each data point represents a single kinetochore. CENP-A levels at each cell cycle stage are normalized to the stage with the highest mean CENP-A levels among all cell cycle stages. The normalized mean CENP-A levels are shown above the dot plot of each cell cycle stage. Error bars represent the standard deviation relative to the mean. Each of the two independent replicates is color-coded. Sample size of each replicate is shown above the dot plot. **c**, Representative images and quantification of CENP-B protein levels at kinetochores throughout the cell cycle.

### Dynamics of CENP-A and CENP-B

CENP-A, a centromere specific histone H3 variant, serves as an essential epigenetic marker for kinetochore assembly^22–24^. Unlike canonical histones, CENP-A is loaded onto centromeres exclusively during early G1 phase^25,26^. Consequently, the pre-existing CENP-A nucleosomes are evenly distributed between the original and newly synthesized centromeric chromatin during S phase. In support of this, our qIF analysis demonstrated that CENP-A levels at kinetochores peaked at G1/S phase and decreased by approximately half in G2 phase, remaining constant until telophase (Fig. 1b). These results underscore the robustness of our qIF technique, allowing for precise measurement of the dynamics of kinetochore proteins throughout the cell cycle.

CENP-B binds a 17-bp CENP-B box sequence in centromeric repetitive DNA via its N-terminal domain (aa 1-125)^27,28^. It contributes to the maintenance of CENP-C levels at centromeres and promotes *de novo* centromere formation^29,30^. Our findings showed that CENP-B levels increased from G1/S phase to G2 phase (G1/S: 0.37; G2: 0.49), likely due to the synthesis of new centromeric DNA (Fig. 1c). Note that CENP-B signal levels from G2 phase to metaphase represents the combination of two pools of CENP-B proteins residing at a pair of sister kinetochores. We also observed a further increase in CENP-B levels during mitotic progression with a peak at metaphase (Fig. 1c). This could be due to the stability of CENP-B at centromeres. While CENP-B exhibits a highly dynamic exchange rate during G1/S phase, it binds stably to centromeres after G2 phase^14^.

### Dynamics of the CCAN

The CCAN provides a structural platform for outer kinetochore assembly, and thereby serves as a bridging module between centromeric chromatin and spindle microtubules. CCAN exhibits high structural flexibility, which is critical for controlling SAC activity and the binding affinity between kinetochores and microtubules^31,32^. Comprising 16 subunits, CCAN is composed of at least five subcomplexes: CENP-C, CENP-T-W-S-X, CENP-N-L, CENP-H-I-K-M, and CENP-O-P-Q-U-R. All CCAN proteins are constitutively present at centromeres throughout the cell cycle, though their kinetic profiles vary^33,34^. To comprehensively analyze the key components of the CCAN complex, we determined the protein dynamics of CENP-C, CENP-N, CENP-I, CENP-K, and CENP-T throughout the cell cycle by qIF (Fig. 2a).

**Fig. 2.**
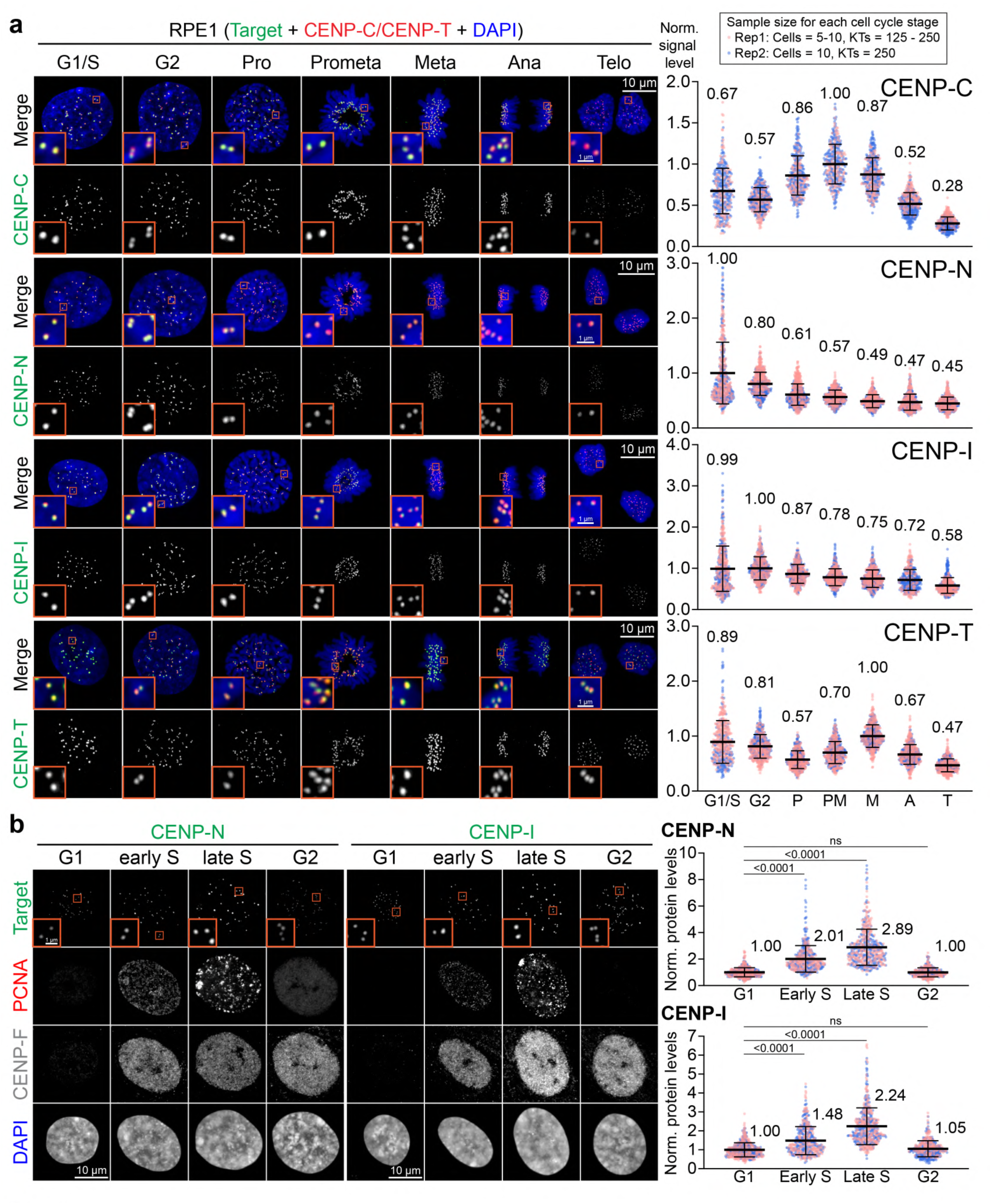
Dynamics of inner kinetochore proteins throughout the cell cycle. **a**, Representative images and quantification of CENP-C, CENP-N, CENP-I, and CENP- T protein levels at kinetochores throughout the cell cycle. **b**, Representative images and quantification of CENP-N and CENP-I protein levels at kinetochores during G1, early S, late S, and G2 phase.

CENP-C directly interacts with CENP-A nucleosomes through its central domain (aa 426-537)^35^ and CENP-C motif (aa 738-758)^36^, enabling one CENP-C molecule to bind to two adjacent CENP-A nucleosomes^37^. While the N-terminal PEST-rich domain (aa 181- 373) allows for the interaction with the CENP-HIKM and CENP-NL subcomplexes^38,39^, the C-terminal Cupin domain (aa 773-943) facilitates CENP-C dimerization and its recruitment to kinetochores^37^. We found that CENP-C levels significantly increased from interphase to prometaphase (G1: 0.67; PM: 1.00), followed by a gradual decline during mitotic progression (M: 0.87; T: 0.28) (Fig. 2a). A similar kinetic pattern was reported using GFP-fused CENP-C along with live-cell imaging in HeLa cells^34^. The variation of CENP-C levels during mitosis cannot be attributed to the abundance of its upstream adaptor, CENP-A, as CENP-A levels remained constant throughout mitosis (Fig. 1b). This variation may instead be influenced by the phosphorylation status of CENP-C by Cdk1, as phosphorylation of CENP-C by Cdk1 has been shown to increase CENP-C’s binding affinity to CENP-A in chicken DT40 cells^36,40^.

CENP-N and CENP-L form a heterodimer through their C-terminal domains^39^. The N-terminal domain of CENP-N (aa 1-286) is responsible for its binding to CENP-A nucleosomes^39,41–43^, while the C-terminus of CENP-N (aa 287-339) associates with CENP-C and the CENP-HIKM subcomplexes, contributing to their kinetochore localization^44^. We found that CENP-N levels peaked during G1/S phase, likely because of the replication of centromeres, and exhibited a gradual, albeit slight decrease as the cell cycle progressed (G1/S: 1.00; G2: 0.80; P-T: 0.61-0.45) (Fig. 2a). The CENP-HIKM subcomplex binds CENP-C, CENP-NL, and CENP-TWSX *in vitro*^44,45^, and these interactions are necessary for its kinetochore localization^39,44^. In line with their reported dependency on kinetochore localization, the kinetics of both CENP-I and CENP-K closely mirrored that of CENP-N (Fig. 2a and Extended Data Fig. 2), displaying a gradual yet steady decline from interphase through the end of mitosis (G1/S: 1.00; T: 0.58 or 0.55) (Fig. 2a and Extended Data Fig. 2). Since CENP-C is essential for the recruitment of CENP-NL and CENP-HIKM, the reduction of CENP-C levels following prometaphase may contribute to the dissociation of these subcomplexes from kinetochores (Fig. 2a).

The CENP-TWSX heterotetramer is formed by the association of a CENP-TW dimer and a CENP-SX dimer^46^. Although CENP-TW and CENP-SX complexes can independently and directly bind to DNA *in vitro*, the CENP-TW complex is required for CENP-SX kinetochore localization in cells^46^. Our qIF showed that CENP-T levels peaked in metaphase (P: 0.57; PM: 0.70; M: 1.00; A: 0.67; T: 0.47) (Fig. 2a). Prior biochemical studies have demonstrated that the CENP-TWSX complex interacts exclusively with the CENP-HIKM subcomplex, but not with CENP-A or other CCAN proteins^44,45,47,48^. Given that levels of CENP-I and CENP-K do not increase from prophase to metaphase (Fig. 2a), the recruitment of additional CENP-T molecules in early mitosis does not primarily depend on the CENP-HIKM subcomplex. The CENP-T homolog in budding yeast, Cnn1, is recruited to kinetochores during mitosis through Cdk1-mediated phosphorylation^49^, suggesting that a similar conserved mechanism may exist in human cells for recruiting additional CENP-T during early mitosis.

We noticed there was a high variance of CCAN signal intensity during G1/S phases compared to other stages of the cell cycle. We hypothesized that, unlike CENP-A, CCAN protein levels varied between G1 and S phases because additional CCAN proteins were immediately assembled on the newly synthesized centromeres during S phase^41,50,51^. To test this hypothesis, we distinguished cells between G1, early S, late S, and G2 phases using the method described in Fig. 1a, and performed qIF for CENP-N and CENP-I. Our findings revealed that both CENP-N and CENP-I levels gradually increased from G1 to early S phase, peaked at late S phase (2-3 fold higher than in G1), and then decreased by half during late G2 phase when sister kinetochore pairs appeared (Fig. 2b). We confirmed that the distance between sister kinetochores on replicated centromeres in S phase remain below the diffraction limit of light microscopy, as evidenced by the consistent number of kinetochore foci from G1 to late S phase (Extended Data Fig. 1b). These results demonstrate that CCAN proteins are immediately recruited to newly synthesized centromeres to form the kinetochores during S phase. Furthermore, we found no significant difference in CENP-N and CENP-I levels between G1 and G2 phases (Fig. 2b), indicating that CCAN proteins are equally distributed between sister kinetochores. In conclusion, CCAN proteins are promptly recruited to newly synthesized kinetochores in nearly equal amounts as the original kinetochores during S phase.

### Dynamics of the KMN network

The KMN network has been believed to be recruited to kinetochores during mitosis^52,53^. Previous studies have demonstrated that the KMN network is assembled at kinetochores through two distinct pathways: the CENP-C and CENP-T pathways^54^. CENP-C recruits an entire KMN network by directly binding to Mis12C through its N-terminus in an Aurora B (AurB)-dependent manner^54–57^. On the other hand, CENP-T can directly recruit up to two Ndc80C and one entire KMN network through Cdk1 phosphorylation^20,58^.

Mis12C, also known as MIND complex in budding yeast, consists of four protein subunits: Mis12, Dsn1, Pmf1, and Nsl1^56,59^. X-ray crystallography has revealed that Mis12C is formed by the intertwined C-terminal segments of four proteins in a 1:1:1:1 stoichiometry^56^. Mis12C is responsible for the linkage between Knl1C and Ndc80C to CENP-C and CENP-T^37,54,56^. Nsl1 mediates the interaction with Knl1^60,61^, and both Dsn1 and Nsl1 are involved in binding to the Spc24-Spc25 subunits of Ndc80C^60^. Our qIF analysis demonstrated that all four Mis12C subunits exhibited similar temporal dynamics (Fig. 3 and Extended Data Fig. 2), indicating that the heterotetrameric Mis12C maintained consistent stoichiometry throughout the cell cycle. Unexpectedly, we found that all Mis12C subunits were detected at kinetochores in G1/S phase (G1/S: 0.14-0.47) (Fig. 3 and Extended Data Fig. 2). Mis12C levels increased and reached a peak during prophase, gradually declined as mitosis progressed, and culminated in mitotic exit (P: 1.00; PM: 0.86-0.54; T: 0.05-0.02). The pronounced increase in Mis12C levels during mitotic entry is attributed to the enhanced kinase activity of AurB and Cdk1, which promotes Mis12C installation on CENP-C and CENP-T, respectively^37,54,55^. The reduction of Mis12C levels at kinetochores during the late stages of mitosis is likely due to the reduced kinase activity of Cdk1 and AurB^62–64^.

**Fig. 3.**
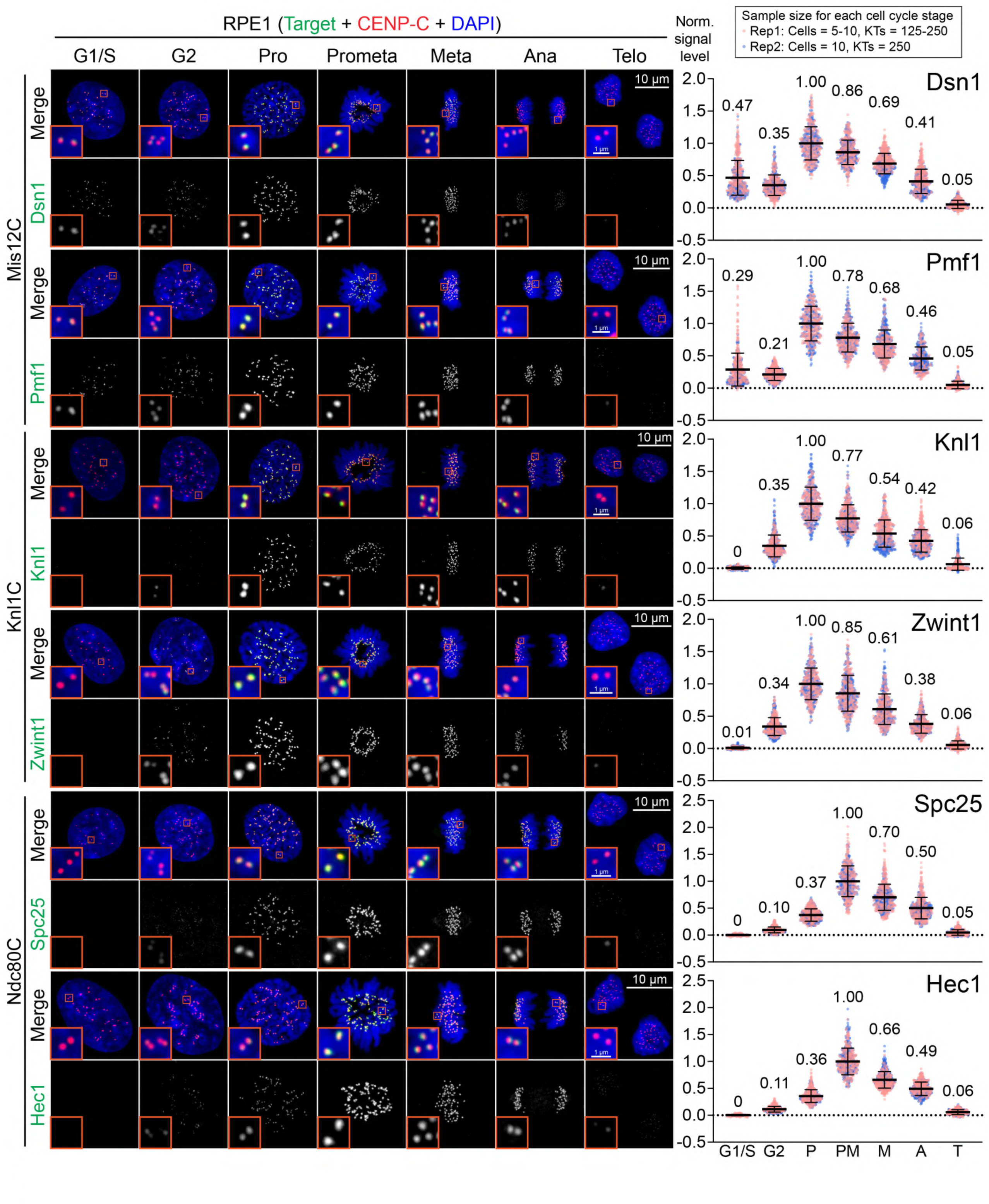
Dynamics of outer kinetochore proteins throughout the cell cycle. Representative images and quantification of Dsn1, Pmf1, Knl1, Zwint1, Spc25, and Hec1 protein levels at kinetochores throughout the cell cycle.

Knl1C is formed by Knl1 and Zwint1 via interactions between Zwint1 and the C-terminal Knl1 (aa 2010-2134)^9,65,66^. Knl1 is essential for the recruitment of Zwint1 to kinetochores, and Zwint1 also partially contributes to the kinetochore localization of Knl1^8,67^. The C-terminus of Knl1 features two RWD domains: RWD^N^ (aa 2109-2209) and RWD^C^ (aa 2210-2311)^61^. Knl1 RWD^N^ binds to Nsl1, whereas RWD^C^ interacts with both Dsn1 and Pmf1^8,9,60,61^. Our qIF data revealed that Knl1 and Zwint1 first appeared at kinetochores in selected cells in late S phase, although not uniformly across all kinetochores. Their signal levels then significantly increased in G2 phase, reaching a peak in prophase (G2: 0.34-0.35; P: 1.00) (Fig. 3 and Extended Data Fig. 3). Following NEBD, levels of both Knl1 and Zwint1 gradually decreased, becoming nearly undetectable by telophase (PM: 0.77-0.85; M: 0.54-0.61; A: 0.38-0.42; T: 0.06). Notably, the kinetic profile of Knl1C mirrored that of its upstream recruiter, Mis12C (Fig. 3). However, Knl1C was completely undetectable at kinetochores from G1 and through middle-to-late S phases, even in the presence of Mis12C. These findings suggest that Knl1C can access interphase nucleus, but additional post-translational modifications (PTMs) may be essential for Knl1’s binding to Mis12C or its nuclear localization.

Ndc80C consists of four proteins: Hec1 (Ndc80), Nuf2, Spc24, and Spc25^68,69^. EM images revealed that Ndc80C forms a long rod with a globular head at each end^69,70^. The N-terminal domains of Hec1 directly binds to microtubules^69^, whereas the C-terminus of Spc24-Spc25 directly interacts with either C-terminal Mis12C or N-terminal CENP-T^8,9,60^. We employed qIF with antibodies targeting Hec1 and Spc25. In line with Knl1C, Ndc80C was initially detected at a small subset of kinetochores in late S phase (Extended Data Fig. 3). Their abundance significantly increased during G2 and prophase, reaching peak levels in prometaphase (G2: 0.07-0.10; P: 0.33-0.37; PM: 1.00) (Fig. 3). Subsequently, their presence diminished as mitosis progressed, becoming nearly undetectable by telophase (M: 0.70-0.74; A: 0.48-0.50; T: 0.05). Like Knl1C, Ndc80C can also translocate into interphase nucleus, through PMTs are likely essential for its localization to Mis12C at kinetochores. The increase of Ndc80C levels from the G2 phase to prometaphase coincides with the increased levels of its upstream recruiters (i.e., Mis12C and CENP-T) and increased Cdk1 activity^71^, which facilitates Ndc80C recruitment in a phosphorylation-dependent manner. In conclusion, our qIF results underscore that the KMN network assembles at kinetochores through a multi-step process. Mis12C is recruited to kinetochores during G1 phase, followed by the recruitment of Knl1C and Ndc80C, beginning in late S phase. Furthermore, Ndc80C levels reaches its peak during prometaphase, while Knl1C levels peaks earlier, during prophase. The delayed recruitment of Knl1C and Ndc80C suggests that Mis12C is not merely an adaptor for these complexes, and additional factors are needed after late S phase to facilitate the binding of Knl1C to Mis12 and Ndc80C to Mis12C and CENP-T.

### Dynamics of Corona/SAC proteins

The SAC serves as a cellular surveillance system that detects erroneous microtubule attachments during mitosis^1,72,73^. The mitotic checkpoint complex (MCC)^74^, comprising Bub1, BubR1, Mad2, and Cdc20, inhibits the anaphase promoting complex/cyclosome (APC/C)^75,76^, an E3 ubiquitination ligase, thereby preventing the proteasomal degradation of Securin and Cyclin B^77^. In the current model of SAC signaling, improper kinetochore-microtubule attachment triggers Mps1-mediated phosphorylation of Knl1, which subsequently facilitates SAC protein recruitment, including Bub3, Bub1 and BubR1^65,78,79^. The N-terminal Knl1 contains 19 repeats of MELT motif^79,80^, which are phosphorylated by Mps1 and Plk1 during early mitosis^81–83^. This phosphorylation enables the docking of Bub1/Bub3 and BubR1/Bub3 complexes^79,80,84^. However, not all MELT motifs exhibit equal affinity for these proteins ^80,85^, resulting in an average of 6-7 Bub1/BubR1 proteins binding to a single Knl1 molecule^85^. BubR1 is critical in recruiting Protein Phosphatase 2A (PP2A)-B56 via its C-terminal KARD domain, which contains three phosphorylation sites for Cdk1 and Plk1^86–88^. The BubR1-bound PP2A-B56 complex subsequently decreases the overall phosphorylation on Knl1 MELT repeats, leading to a marginal reduction in Bub1 levels at kinetochores^83,86,89,90^. Upon chromosome biorientation, PP1 phosphatase is recruited to kinetochores, which further extinguishes SAC activity^83,89,91^.

Bub1 recruitment to kinetochores began during G2 phase and peaked in prophase (G2: 0.05; P: 1.00) (Fig. 4), matching the kinetics of Knl1 phosphorylation at MELT motifs (hereafter termed pMELT) (G2: 0.01; P: 1.00) (Fig. 4). Notably, BubR1 recruitment exhibited a slight but significant delay, appearing at kinetochores from late prophase (Fig. 4 and Extended Data Fig. 4). After NEBD, Bub1 levels decreased slightly (PM: 0.72), while BubR1 levels peaked (PM: 1.00). During metaphase, both Bub1 and BubR1 levels underwent a dramatic reduction (M: 0.13-0.25), consistent with the reduction of pMELT levels (PM: 0.45; M: 0.03). Interestingly, BubR1-S670 is highly phosphorylated during late prophase and prometaphase (PM: 1.00), suggesting immediate phosphorylation by Cdk1 upon its kinetochore recruitment. This phosphorylation event likely promotes PP2A-B56 recruitment, leading to a moderate suppression of SAC activity, as evidenced by the reduction in pMELT and Bub1 levels during prometaphase (Fig. 4). We observed that a small fraction of Bub1 and BubR1 persisted at kinetochores from metaphase to anaphase (M: 0.13-0.25; A: 0.07-0.14), even though the pMELT levels became almost undetectable by metaphase (M: 0.03) (Fig. 4). This observation suggests the presence of two distinct pools of these proteins: one that is stably associated with kinetochores, independent of MELT phosphorylation status, and another whose binding to Knl1 is dependent on SAC activity.

**Fig. 4.**
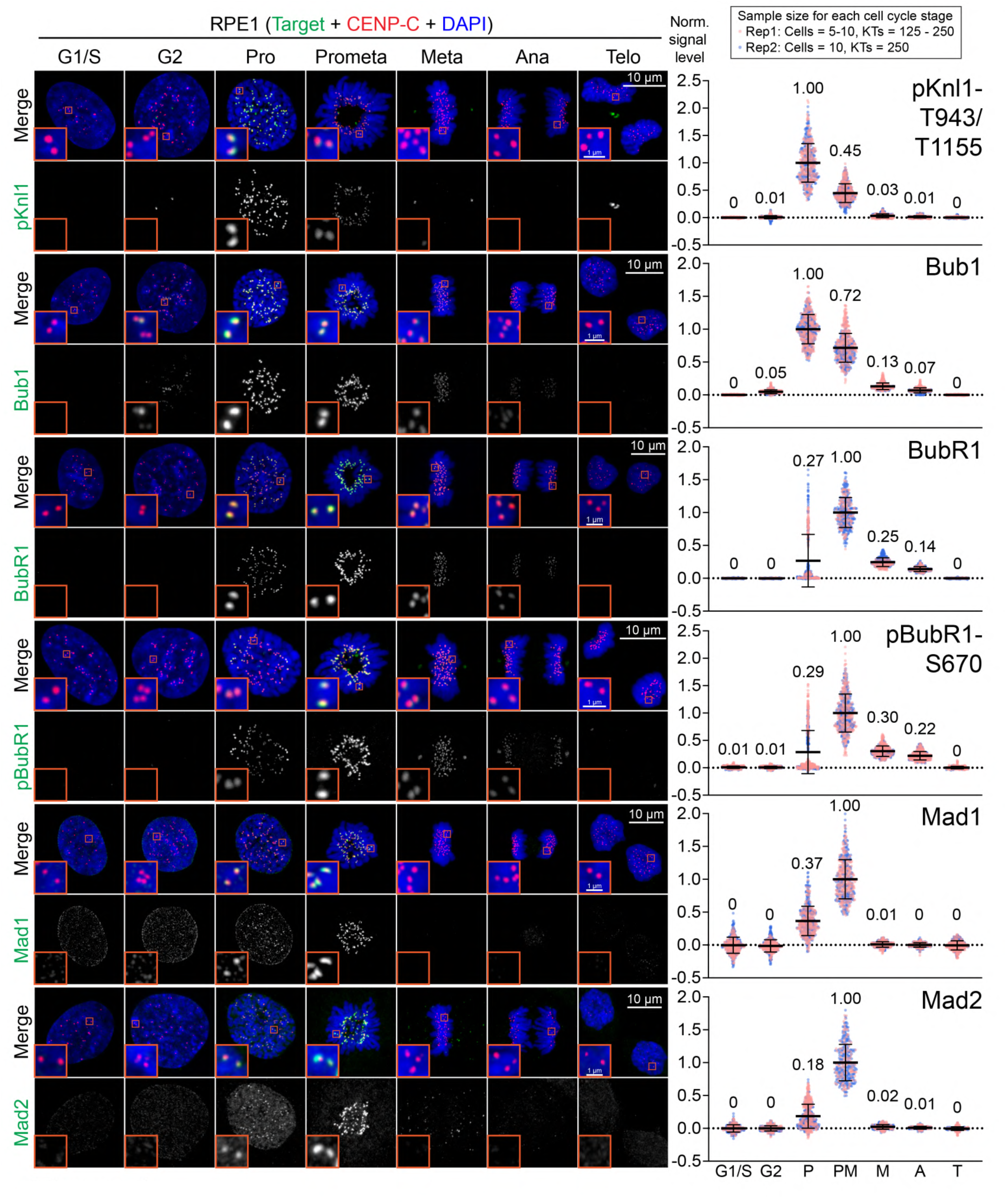
Dynamics of spindle checkpoint proteins and SAC-related phosphosites throughout the cell cycle. Representative images and quantification of pKnl1-T944/T1155, Bub1, BubR1, pBubR1- S670, Mad1, and Mad2 protein levels at kinetochores throughout the cell cycle.

Kinetochore localization of Bub3, a critical binding partner of Bub1 and BubR1, displayed consistent kinetics. Bub3 levels peaked and remained from prophase through prometaphase, followed by a significant reduction during metaphase, becoming undetectable by anaphase (P: 0.95; PM: 1.00; M: 0.16; A: 0.02) (Extended Data Fig. 2). Notably, the Bub3 binding sites on Knl1 (pMELT) were reduced by half from prophase to prometaphase (P; 1.00; PM: 0.45) (Fig. 4), whereas Bub3 levels remains stable (P: 0.95; PM: 1.00) (Fig. 4). This constancy may be attributed to the dissociation of the Bub1-Bub3 complex from prophase to prometaphase (P: 1.00; PM: 0.72) along with the increased recruitment of the BubR1-Bub3 complex to kinetochores (P: 0.27; PM: 1.00), thereby maintaining a constant total copy number of Bub3 at Knl1 during these phases.

Mad2 exists in two distinct conformations: open (O-Mad2) and closed (C-Mad2)^92^. Upon binding to Mad1, Mad2 undergoes a conformational change from its open to closed form^93,94^. The formation of the Mad1/C-Mad2 complex at kinetochores creates a platform that facilitates the further recruitment of cytosolic O-Mad2, which is subsequently converted to C-Mad2^92,95^. We revealed that both Mad1 and Mad2 exhibited similar dynamic profiles at kinetochores, with their recruitment initiating in prophase and reaching their peak in prometaphase (P: 0.18-0.37; PM: 1.00) (Fig. 4). Both Mad1 and Mad2 signals at kinetochores immediately became undetectable in metaphase (M: 0.01-0.02). Mad1 directly binds to Bub1 through phosphorylation by Cdk1 and Mps1^96–99^. Besides Bub1, the RZZ complex has been proposed as an upstream recruiter of Mad1 at kinetochores^100–103^, however, the direct binding between Mad1 and the RZZ complex has yet to be demonstrated. Collectively, our analysis demonstrated that Mad1 and Mad2 were specifically recruited to kinetochores during prophase and prometaphase (Fig. 4). Additionally, consistent with previous observation, both Mad1 and Mad2 were detected on the nuclear pores (NPs) during interphase (Fig. 4 and Extended Data Fig. 5)^104–106^.

The Rod-Zwilch-Zw10 (RZZ) complex assembles as a hexamer with a 2:2:2 stoichiometry in an antiparallel configuration^107^. This complex is crucial for SAC silencing by recruiting Spindly, the adaptor for Dynein^108,109^. Dynein, a minus-end directed motor protein, inactivates SAC by stripping SAC proteins from kinetochores upon proper microtubule attachment^110^. Recent studies have shown that the RZZ complex can self-oligomerize to form a higher-order structure known as fibrous corona^107,111^. Zwint1 is essential for recruiting RZZ complex to kinetochores through a direct interaction with N-terminal Zw10 (aa 1-80), which is regulated by AurB phosphorylation^67,112–114^. Our qIF demonstrated that both Rod and Zw10 levels peaked during prometaphase, decreased substantially in metaphase, and became barely detectable in anaphase (PM: 1.00; M: 0.21; A: 0.05) (Fig. 5). We noticed that both Zwint1 and AurB were present at kinetochores during G2 phase (Fig. 3 and 6), whereas RZZ complex did not localize to kinetochores until prometaphase, possibly due to the absence of nuclear localization signals (NLS) in RZZ proteins^115^.

**Fig. 5.**
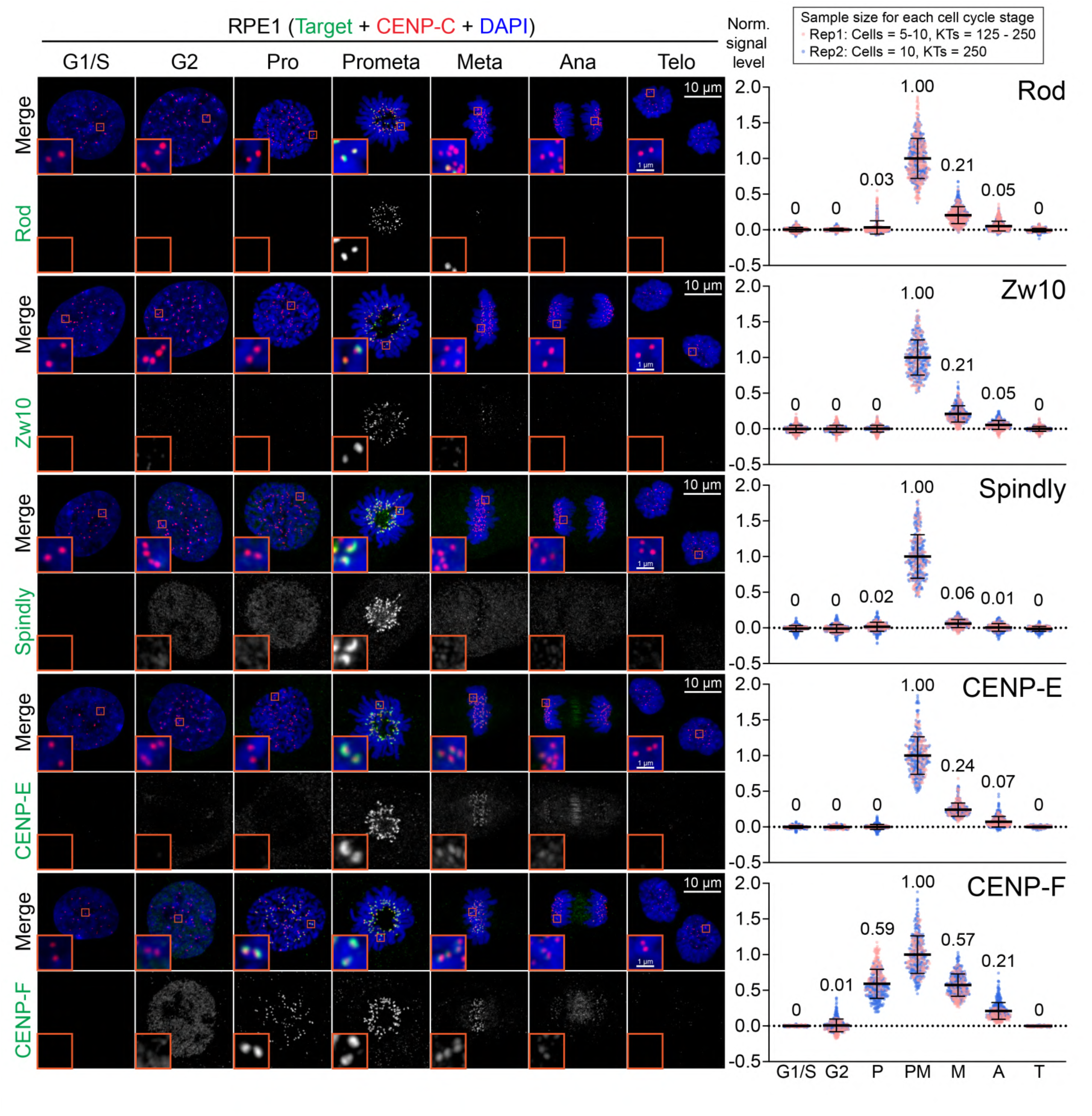
Dynamics of corona proteins throughout the cell cycle. Representative images and quantification of Rod, Zw10, Spindly, CENP-E, and CENP-F protein levels at kinetochores throughout the cell cycle.

**Fig. 6.**
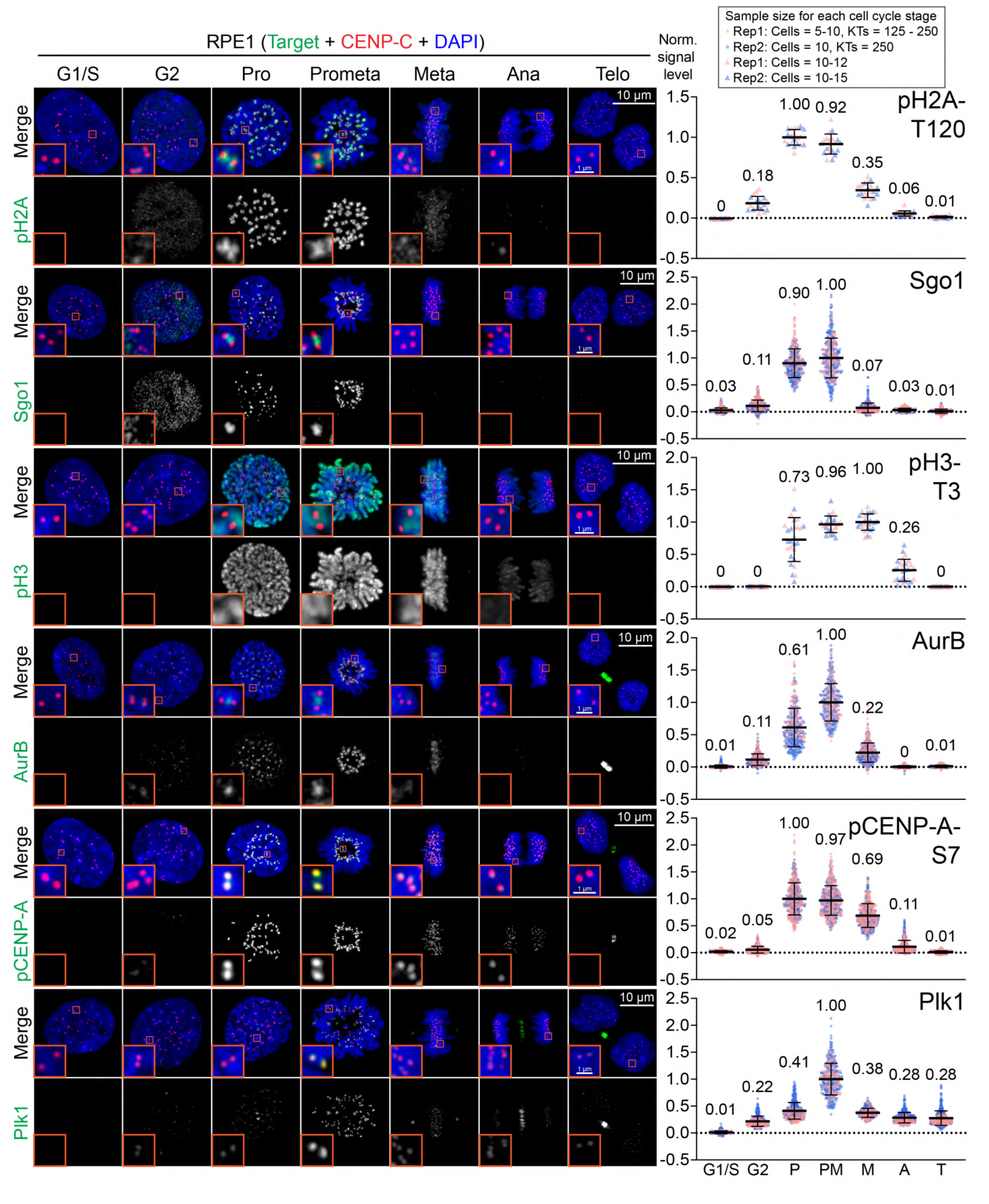
Dynamics of kinases and kinase-related phosphosites throughout the cell cycle. Representative images and quantification of pH2A-T120, Sgo1, pH3-T3, AurB, pCENP- A-S7, and Plk1 protein levels at kinetochores throughout the cell cycle.

Spindly, recruited by the RZZ complex, plays dual roles at kinetochores: it promotes corona assembly^107^ and serves as an adaptor for Dynein, thereby contributing to the inactivation of SAC^110,116,117^. Consistent with its interaction with the RZZ complex, Spindly predominantly localized to kinetochores during prometaphase followed by a significant loss in metaphase (PM: 1.00; M: 0.06) (Fig. 5). Except for these two mitotic stages, Spindly signals were absent from kinetochores, but it demonstrated distinct nuclear accumulation during G2 and prophase (Fig. 5 and Extended Data Fig. 5).

CENP-E, a plus-end directed kinesin-like motor protein, promotes chromosome congression and facilitates the transition from lateral to end-on microtubule attachment^118–120^. Bub1, BubR1, Mad1, and the RZZ complex have been proposed as upstream recruiters of CENP-E^121–124^, however, further investigation is required to clarify their specific roles in this process^124–126^. During prometaphase, CENP-E was robustly recruited to kinetochores, reaching its peak levels (PM: 1.00) (Fig. 5). CENP-E levels significantly decreased in metaphase and continued to decline through anaphase (M: 0.24; A: 0.07). This kinetic profile closely matched that of the RZZ complex but differed from those of Bub1, BubR1, and Mad1/2 (Fig. 4 and 5), which were recruited to kinetochores prior to NEBD. Bioinformatic analysis indicates that CENP-E lacks NLS^115^, suggesting that NEBD may be required for its targeting to kinetochores.

Unlike CENP-E, CENP-F lacks a motor domain but contains two microtubule-binding domains^127,128^. CENP-F is recruited to kinetochores through its interaction with the kinase domain of Bub1, a process regulated by AurB activity^123,125^. CENP-F began localizing to kinetochores during prophase, reaching its peak in prometaphase (P: 0.59; PM: 1.00) (Fig. 5). CENP-F levels then gradually declined from metaphase to anaphase and eventually became undetectable during telophase (M: 0.57; A: 0.21; T: 0). We observed that ti50% of CENP-F remained at kinetochores during metaphase (Fig. 5), while Bub1 levels markedly reduced (M: 0.13) (Fig. 4). This suggests that Bub1 is either not essential for the retention of CENP-F at kinetochores or that only a small fraction of Bub1 associates with CENP-F in prometaphase. Despite the absence of CENP-F at kinetochores, CENP-F accumulated in the nucleoplasm from S phase to G2 phase (Fig. 1a and Extended Data Fig. 1a). The lack of CENP-F at kinetochores during these stages indicates that CENP-F alone is insufficient for its kinetochore targeting, and likely Bub1 and AurB activity are also required^125,129^. Collectively, our findings reveal a subtle yet significant temporal gaps in the recruitment and dissociation of SAC-related proteins at kinetochores.

### Dynamics of mitotic kinases

Mitotic kinases orchestrate kinetochore functions by controlling the assembly of kinetochore proteins and modulating the binding affinity between kinetochores and microtubules. Understanding the temporal dynamics of these kinases helps us unravel the sequential recruitment events of kinetochore proteins during mitosis. Therefore, we directly or indirectly examined the dynamics of five major mitotic kinases throughout the cell cycle: Sgo1, Sgo2, AurB, Plk1, and Haspin.

The linkage between sister chromatids is established during S phase by the cohesin complex^130–132^. Upon mitotic entry, most cohesin complexes are removed from chromosomes, except for centromeres, where they are protected by Sgo1^133,134^. Sgo1 is recruited to kinetochores via Bub1-dependent phosphorylation of histone H2A at Thr120 (pH2A-T120)^135–137^. Our qIF demonstrated that Sgo1 began accumulating in the nucleus, including, but not limited to, the centromere regions, during the G2 phase (G2: 0.11) (Fig. 6). This initial nuclear accumulation of Sgo1 coincides with the appearance of pH2A-T120 signals throughout the nucleus (Fig. 6 and Extended Data Fig. 5), suggesting that Sgo1 can immediately associate to H2A histones upon phosphorylation by Bub1. The levels of centromere-bound Sgo1 increased and peaked during prophase and prometaphase (P: 0.90; PM: 1.00). Subsequently, Sgo1 levels decreased rapidly to near-background levels during metaphase (M: 0.07). This substantial loss of Sgo1 is attributed to the marked reduction in pH2A-T120 levels (PM: 0.92; M: 0.35) (Fig. 6), which is likely driven by the recruitment of PP1 and PP2A.

Sgo2 plays a crucial role in protecting chromatid cohesion from premature cleavage during meiosis^138,139^. In mitosis, Sgo2 localizes to inner centromeres, though its precise function and recruitment mechanism remain largely unexplored. Like Sgo1, Sgo2 began accumulating at centromeres during G2, peaked in prophase, and gradually decreased, becoming nearly undetectable by anaphase (G2: 0.10; P: 1.00; PM: 0.83; M: 0.48; A: 0.03) (Extended Data Fig. 2). Interestingly, unlike Sgo1, ∼50% of Sgo2 remained associated with centromeres during metaphase. This suggests that additional factors contribute for its retention at centromeres, consistent with its partial recruitment dependence on Mps1 and Bub1^140^.

The Chromosomal Passenger Complex (CPC) consists of four subunits: AurB kinase, INCENP, Borealin, and Survivin^141^. INCENP serves as a scaffold linking the AurB kinase and the localization modules (Borealin and Survivin)^141,142^. CPC localizes to the inner centromeres and kinetochores via two distinct pathways^143,144^. In the first pathway, Haspin kinase phosphorylates histone H3 at Thr3 (pH3-T3), facilitating Survivin binding^143,145,146^. In the second pathway, Bub1 kinase phosphorylates H2A-T120, which enables Sgo1 binding and the subsequent recruitment of Borealin^143,147^. During anaphase, CPC is translocated to the spindle midzone, and in telophase, it relocates to the midbody^148–151^. We demonstrated that AurB began localizing to kinetochores during G2 phase (G2: 0.11) (Fig. 6 and Extended Data Fig. 5), coinciding with the initial association of Sgo1 with histones (G2: 0.11) (Fig. 6). Concurrently, punctate signals of pH3-T3 were detected outside centromeres in the G2 nucleus, but these levels were relatively low compared to its maximum levels (G2: 0) (Fig. 6 and Extended Data Fig. 5). Like Sgo1 and pH3-T3, AurB levels substantially increased from prophase to prometaphase (P: 0.61; PM: 1.00). This increase aligns with AurB’s role in destabilizing improper kinetochore-microtubule attachments, facilitating error correction. During metaphase, both AurB and Sgo1 showed a marked decrease in their kinetochore levels, though they did not completely fall to background levels (AurB-M: 0.22; Sgo1-M: 0.07). In contrast, pH3-T3 levels remained high during this stage (M: 1.00). Notably, AurB levels at the midzone during anaphase were significantly lower compared to its centromere levels from prophase to metaphase and at the midbody during telophase (Fig. 6). To further assess AurB kinase activity, we examined its downstream substrate CENP-A Ser7 (CENP-A- S7)^152–154^. Phosphorylation of CENP-A-S7 was detected from G2 phase, peaked during prophase, and sustained through prometaphase (G2: 0.05; P: 1.00; PM: 0.97) (Fig. 6). Notably, the peak of pCENP-A-S7 occurs earlier than that of AurB, suggesting that complete AurB loading to centromeres is not essential for reaching maximal pCENP-A- S7 levels. The pCENP-A-S7 levels gradually declined from metaphase to anaphase (M: 0.69; A: 0.11), likely due to the dissociation of AurB and the recruitment of phosphatases.

Plk1, a serine/threonine protein kinase, governs multiple essential processes throughout mitosis, including centrosome separation and maturation, chromatin condensation, kinetochore-microtubule attachment, spindle assembly, and cytokinesis^155^. Its dynamic subcellular localization enables interactions with various substrates^156,157^. The recruitment of Plk1 to kinetochores is proposed to be mediated by Bub1 and CENP- U in a Cdk1-dependent manner^158,159^. Plk1 was present at kinetochores from G2 phase through telophase with its peak levels in prometaphase (G2: 0.22; PM: 1.00; T: 0.28) (Fig. 6). Notably, although both Bub1 and Plk1 began localizing to kinetochores during G2 phase, Plk1 demonstrated a significant delay in reaching its peak, implying the requirement for additional PTMs for its full recruitment. Consistent with previous studies^157,160^, Plk1 was also detected at centrosomes from G2 to telophase, at the central spindle in anaphase, and at the midbody in telophase (Fig. 6).

### The Ska complex detects microtubule attachments, while Astrin-SKAP acts as a tension sensor

The Ska complex is considered a functional homologue of the yeast Dam1 complex, which is responsible for stable kinetochore-microtubule interactions^161,162^. It comprises three proteins: Ska1, Ska2, and Ska3^163^, all of which exhibit interdependence for their recruitment to kinetochores^162,164^. Prior studies have demonstrated that the presence of Ndc80C and the inhibition of AurB activity are critical for Ska complex recruitment to kinetochores^164–166^. As anticipated, Ska3 recruitment to kinetochores began in early prometaphase following NEBD (Fig. 7a), indicating that most kinetochores at this stage are already microtubule-bound. Notably, Ska3 levels were saturated from prometaphase to metaphase (PM: 1.00; M: 0.96) (Fig. 7a), suggesting that its recruitment is a binary, all-or-none process. Ska3 levels then decreased during anaphase and became undetectable by telophase (A: 0.58; T: 0.09), likely correlating with a reduction in Ndc80C (Fig. 3). Besides its kinetochore localization, Ska3 was also detected at the spindle poles from prophase to anaphase (Fig. 7a), aligning with previous research ^162,164,167^. In summary, our qIF analysis supports the model in which Ndc80C and microtubule attachments are key drivers for Ska complex recruitment to kinetochores during unperturbed mitosis. However, while the Ska complex is indicative of kinetochores-microtubule attachment, it does not differentiate between lateral or end-on attachment modes.

**Fig. 7.**
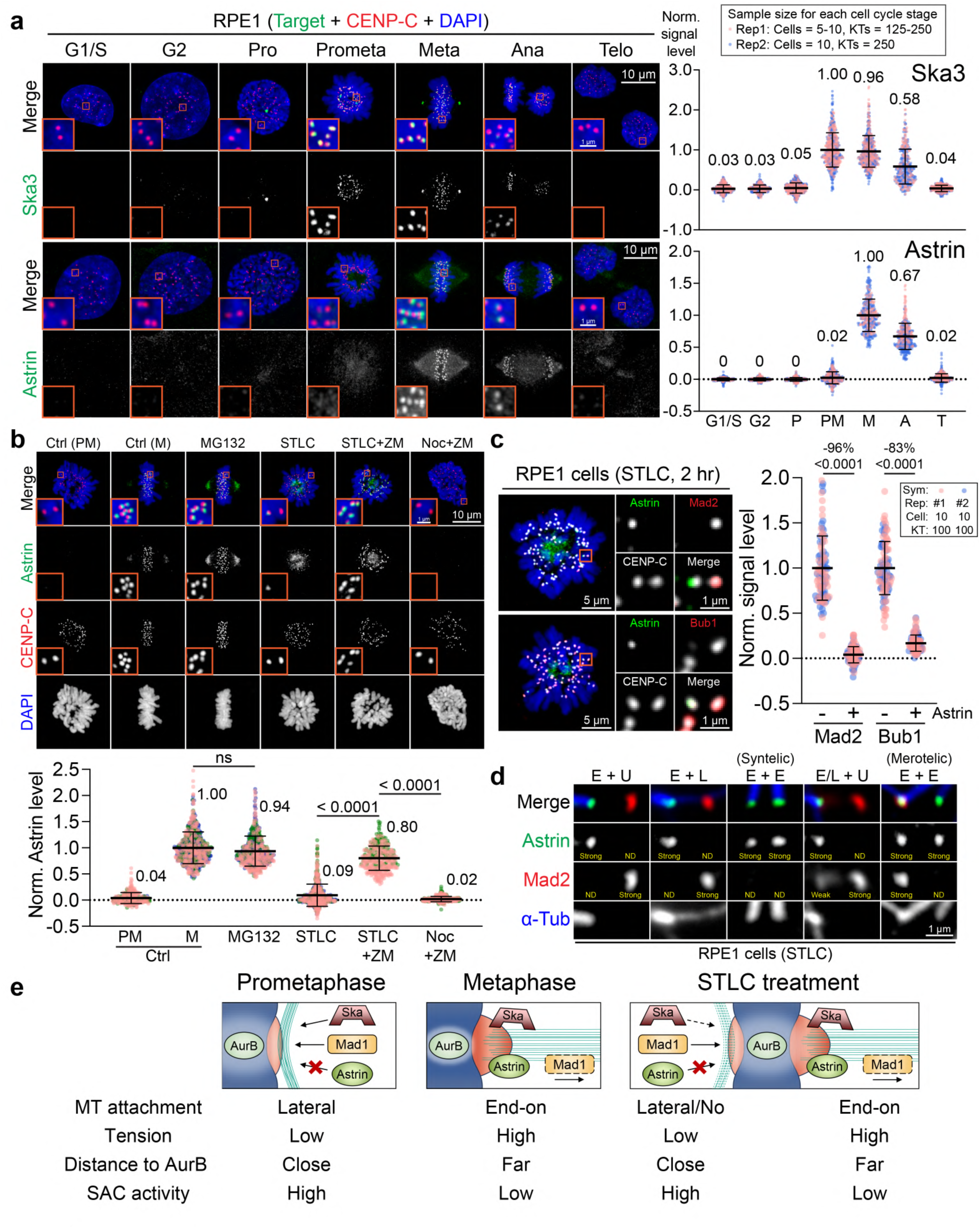
Recruitment of Astrin and Ska complex to kinetochores. **a**, Representative images and quantification of Ska3 and Astrin protein levels at kinetochores throughout the cell cycle. **b**, Top: representative images of untreated control RPE1 cells or cells treated with 10 μM MG132, 2.5 μM STLC, 2.5 μM STLC and 10 μM ZM447439, or 3 μM Nocodazole and 10 μM ZM447439 for 2 hours and stained with Astrin. Bottom: quantification of Astrin levels at kinetochores in each condition. Each data point is a single kinetochore. Three independent replicates were performed and color-coded for each condition. The mean value is shown at the top right of each dot plot. The p-value was calculated using Tukey’s multiple comparisons test. **c**, Left: representative images of RPE1 cells treated with 2.5 μM STLC for 2 hours and stained with indicated antibodies and DAPI. An example pair of sister kinetochores, located in the orange box, is shown in an enlarged image. Right: quantification of Mad2 and Bub1 signal levels at Astrin-positive and Astrin-negative kinetochores within the same sister kinetochore pair. Each data point is a single kinetochore. Two independent replicates were performed and color-coded. The p-value was calculated using Student’s t-test. **d**, Representative images of Astrin, Mad2, and microtubules in RPE1 cells treated with 5 μM STLC for 2 hours. The relative signal intensity of Astrin and Mad2 within the sister kinetochore pair is labeled in yellow. E: end-on attachment; L: lateral attachment; U: unattached; ND: Non-detected. **e**, Schematic diagram of the relationship between microtubule binding status, kinetochore tension, kinetochore stretching, SAC activity, and the recruitment of kinetochore proteins (including Ska complex, Mad1, and Astrin) in unperturbed normal prometaphase cells, normal metaphase cells and STLC-treated cells with monopolar spindles.

The Astrin-SKAP complex (hereafter termed Astrin-SKAP) localizes to both kinetochores and mitotic spindles during mitosis due to its capacity to directly interact with Ndc80C and tubulin^168–171^. The localization of Astrin and SKAP at kinetochores exhibit a reciprocal dependency^170^. The key function of Astrin-SKAP is to stabilize kinetochore-microtubule attachments, thereby facilitating chromosome congression^171–173^. Similar to the Ska complex, Astrin-SKAP recruitment to kinetochores requires the presence of Ndc80C and the inhibition of AurB kinase activity^168,170^. Strikingly, our qIF revealed that Astrin signals were undetectable at kinetochores during early prometaphase (PM: 0.02) (Fig. 7a). Its kinetochore localization suddenly began and reached the peak levels in metaphase (M: 1.00). Astrin levels then moderately decreased during anaphase and became nearly undetectable during telophase (A: 0.67; T: 0.02).

Our qIF analysis revealed that both Mad1/Mad2 and Ska3 reached their peak levels during early prometaphase (Fig. 4 and 7a). This observation suggests that, at early prometaphase, most kinetochores are associated with microtubules through either lateral or end-on attachments, but the tension is still insufficient to eliminate SAC proteins from kinetochores. To further dissect the mechanism of Astrin-SKAP recruitment to kinetochores, we examined Astrin levels under the following conditions. First, we investigated whether increased tension could promote additional Astrin recruitment. To this end, Astrin levels were assessed in cells treated with MG132, a proteasome inhibitor known to enhance kinetochore tension^31^. Unexpectedly, Astrin levels in MG132-treated cells were comparable to those in control metaphase cells (Fig. 7b), indicating that Astrin levels were saturated in normal metaphase cells.

Next, we explored the necessity of kinetochore biorientation for Astrin recruitment by using STLC, an Eg5 inhibitor that induces monopolar spindles^174^. We found that Astrin only localized to a small subset of kinetochores after 2 hours of treatment with STLC (Fig. 7b). Additionally, those Astrin-positive kinetochores were the ones close to the center of monopolar spindle within a pair of sister kinetochores (Fig. 7b). Aligned with the previous research^165^, simultaneous treatment with STLC and ZM447439, an AurB inhibitor, resulted in nearly uniform Astrin levels across all kinetochores with comparable levels to those in control metaphase cells, whereas Astrin signals were completely absent from kinetochores in the presence of both nocodazole and ZM447439 (Fig. 7b). Combined with the qIF data of Ska3, these findings suggest that the AurB activity in early prometaphase kinetochores is still too high to recruit Astrin even though kinetochores are attached by microtubules and potentially generating tension. To further understand the relationship between Astrin recruitment and the SAC activity on the same kinetochore, we co-stained Astrin with SAC proteins, including Mad2 and Bub1. Under the same condition, Astrin-positive kinetochores exhibited negligible Mad2 signals and significantly decreased Bub1 levels compared to Astrin-negative kinetochores (Fig. 7c), indicating that Astrin preferentially localizes to kinetochores with diminished SAC activity. Note that the remaining Bub1 levels in Astrin-positive kinetochores in this experiment (17%) (Fig. 7c) were similar to those in untreated metaphase cells (M: 0.13) (Fig. 4), indicating that sister kinetochores with Astrin signals in this condition resemble bioriented metaphase kinetochores.

To ascertain the microtubule binding status under these conditions, cells were incubated in cold media for 10 min before fixation (cold-stability assay), followed by co-staining for Astrin, Mad2, and microtubules. As expected, robust Astrin signals were detected on kinetochores with end-on attachments, devoid of Mad2 signals (Fig. 7d). Kinetochores with lateral attachments showed strong Mad2 signals and no Astrin signals (Fig. 7d). Kinetochores exhibiting both Astrin and Mad2 signals were rarely observed, however, these kinetochores were prone to attachment errors, particularly in instances of merotelic attachments (Fig. 7d). In conclusion, while kinetochore biorientation is not essential for Astrin recruitment, the formation of end-on attachments, including syntelic and merotelic, and the generation of force that locally reduces AurB activity within a single kinetochore are crucial. Furthermore, although both the Ska complex and Astrin-SKAP require low local AurB activity for their kinetochore localization, the Ska complex is more sensitive to the subtle decrease of AurB activity than Astrin-SKAP, as only Ska3 is detected in early prometaphase kinetochores (Fig. 7a).

### Most kinetochores attach to microtubules in early prometaphase, but generate minimal tension

Our comprehensive analysis of kinetochore protein dynamics provides deeper insights into the relationship between microtubule attachment status and its downstream molecular responses. Our qIF of the SAC, Ska complex, and Astrin-SKAP revealed that most kinetochores are laterally attached to microtubules in early prometaphase, when chromosomes are arranged as a rosette (Fig. 7e and Extended Data Fig. 6). While these lateral attachments efficiently recruit both SAC proteins and the Ska complex, they fail to recruit Astrin-SKAP (Fig. 7e). This suggests that the tension exerted on kinetochores at this stage is insufficient to reduce AurB activity for Astrin-SKAP recruitment and to remove SAC proteins. In other words, microtubule attachment alone is insufficient to inactivate the SAC, and proper tension is required. Consequently, the recruitment of Astrin-SKAP and the retention of SAC proteins at kinetochores are nearly mutually-exclusive events. Supporting this, in cells treated with STLC, a subset of kinetochores within sister pairs can generate force through end-on attachments without achieving biorientation (Fig. 7b,d). These kinetochores possess Astrin-SKAP and lack SAC proteins. This evidence underscores the critical role of the Ska complex as a marker for kinetochores engaged in microtubule attachment regardless of lateral or end-on. In contrast, the Astrin-SKAP complex signifies kinetochores under tension, albeit its detection of tension levels is constrained by a relatively narrow dynamic range.

### CENP-C functions as the primary adaptor for recruiting the Mis12C to interphase kinetochores

We demonstrated that Mis12C is recruited to kinetochores as early as the G1 phase, distinguishing its dynamics from other KMN network components (Fig. 3). Approximately 80% of G1 and 100% of S phase cells were Dsn1/Pmf1-positive (Extended Data Fig. 7a,b). Notably, the subset of G1 cells lacking Mis12C corresponded to early G1 cells, immediately following cytokinesis (Extended Data Fig. 7c, 8a-c). Furthermore, Mis12C was also recruited in interphase HeLa cells, suggesting a conserved mechanism across cell types (Extended Data Fig. 7a,b). To investigate whether Mis12C forms the same heterotetrametric complex at interphase kinetochores as observed during mitosis, we quantified the population of interphase cells positive for Dsn1 and Pmf1 as well as the signal intensities for these proteins in Dsn1-AID RPE1 cells^175^. Following 9 hours of Auxin treatment, Dsn1-AID cells exhibited a significant reduction (ti90%) in the percentage of Dsn1- and Pmf1-positive interphase cells, with a nearly complete loss (∼100%) of Dsn1 and Pmf1 signals at kinetochores (Fig. 8a). These findings indicate that Mis12C likely assembles into a heterotetramer complex at interphase kinetochores, akin to its mitotic organization.

**Fig. 8.**
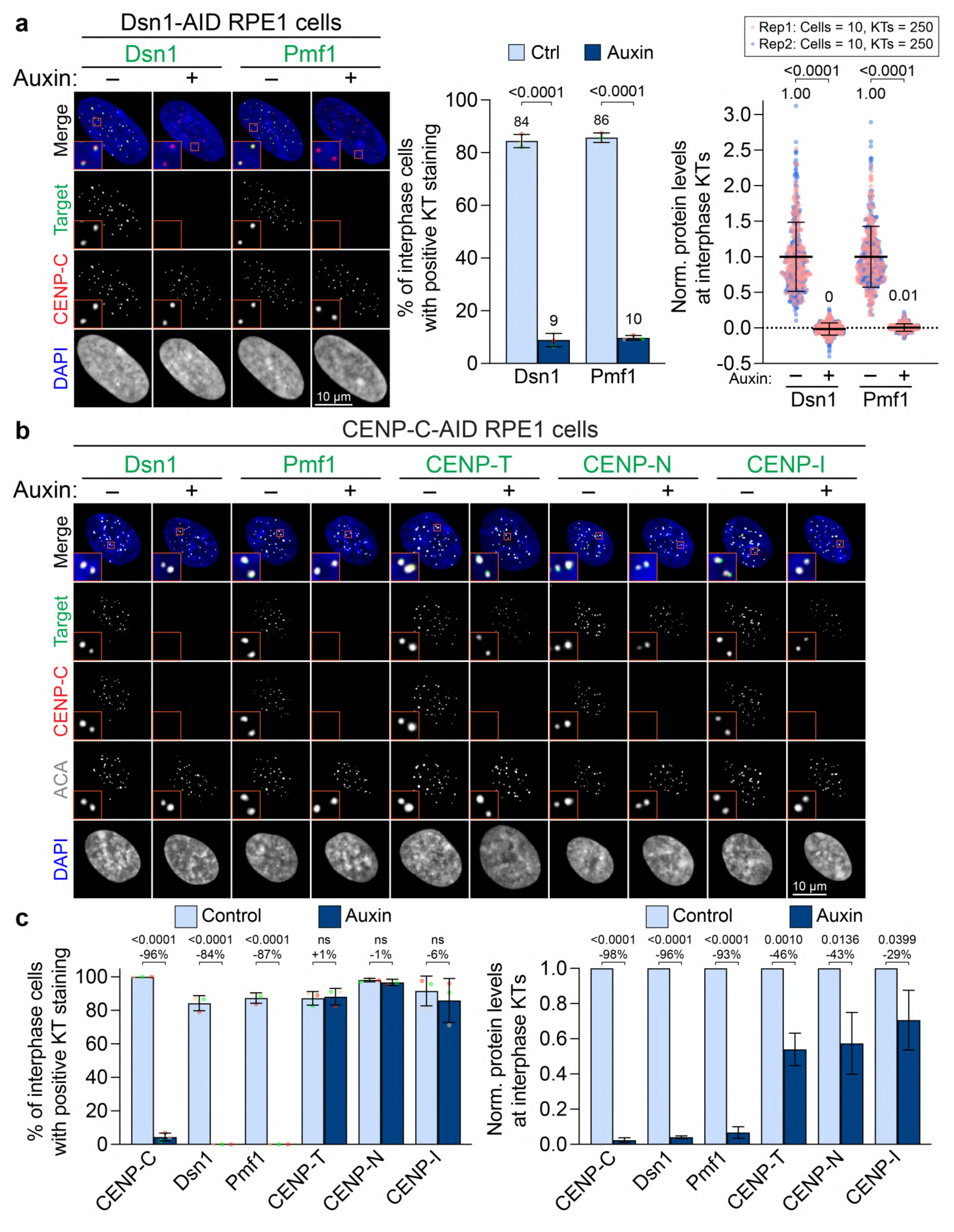
Recruitment of the Mis12C to kinetochores during interphase. **a**, Left: representative images of Dsn1-AID RPE1 cells treated with or without Auxin for 9 hours and stained with the indicated antibodies and DAPI. Middle: quantification of the frequency of interphase RPE1 cells showing Dsn1/Pmf1 kinetochore staining. Each data point is an independent experiment. Three independent replicates were performed and color-coded. Results are the mean ± s.d. The p-value was calculated using Student’s t-test. Right: the normalized signal levels of Dsn1/Pmf1 at kinetochores in untreated control and Auxin-treated interphase cells. Each data point is a single kinetochore. Two independent replicates were performed and color-coded. The p-value was calculated by Student’s t-test. **b**, Representative images of CENP-C-AID RPE1 cells treated with or without Auxin for 1 hour and stained with the indicated antibodies and DAPI. **c**, Left: quantification of the frequency of interphase RPE1 cells showing positive kinetochore staining of indicated proteins. Each data point is an independent experiment. Three independent experiments were performed and color-coded. Results are the mean ± s.d. The p-value was calculated using Student’s t-test. Right: quantification of the normalized indicated protein levels at kinetochores. Results are the mean ± s.d. In each independent experiment, 250 kinetochores from 10 cells were analyzed. The p-value was calculated using Student’s t-test.

Given that both CENP-C and CENP-T independently contribute to Mis12C recruitment to kinetochores during mitosis^54^, we sought to investigate their roles in Mis12C recruitment during interphase. To this end, we utilized a CENP-C-AID RPE1 cell line^36^ to quantify the population of interphase cells positive for Dsn1 and Pmf1, as well as the signal intensity of these proteins at kinetochores upon CENP-C depletion (Fig. 8b,c). Depletion of CENP-C by an hour of Auxin treatment resulted in an 85% reduction in the percentage of Dsn1- and Pmf1-positive interphase cells and a 95% loss of their kinetochore levels (Fig. 8b,c). Since CENP-C stabilizes other CCAN proteins^38,44^, we also quantified the levels of additional CCAN components in CENP-C depleted cells. Notably, we observed no reduction in the percentage of CENP-T, CENP-N, or CENP-I-positive cells upon CENP-C depletion, however, their kinetochore signals were reduced by approximately 50% compared to control interphase cells (Fig. 8b,c). These results indicate that CENP-C, rather than CENP-T or other CCAN proteins, serves as the primary adaptor for Mis12C recruitment to kinetochores in interphase. As Mis12C binding to CENP-C during mitosis is regulated by AurB kinase activity^54–56^, which is thought to be minimal in interphase, we next investigated whether AurB kinase activity is required for Mis12C localization to kinetochores in interphase. To address this, we quantified Dsn1 levels at kinetochores during S phase and prometaphase in RPE1 treated with a high concentration of ZM447439 (Extended Data Fig. 9a). After one hour of treatment, AurB activity was completely abolished, as evidenced by the complete loss of pH3-S10 signals in prometaphase cells (Extended Data Fig. 9b,c). Consistent with previous studies^54,55^, ZM447439 treatment led to a 43% reduction of Dsn1 levels at prometaphase kinetochores (Extended Data Fig. 9c). However, no reduction in Dsn1 levels was observed at interphase kinetochores (Extended Data Fig. 9d), indicating that the mechanism of Mis12C recruitment by CENP-C during interphase is distinct from that in mitosis and independent of AurB kinase activity.

### Limitations of the study

In this study, to quantify protein levels at kinetochores, we used antibodies for fluorescent labeling. However, the binding affinity of antibodies to target proteins or potential conformational changes of the target proteins can affect staining quality. To minimize this, we optimized staining protocols for each antibody, including fixation methods and working concentrations, to achieve a high S/N ratio. We also quantified multiple components within the same protein complex to ensure accurate measurements. While fluorescent proteins (FPs) are valuable tools in studying protein dynamics in live cells, qIF offers distinct advantages, including high detection sensitivity and technical feasibility. Reproducing similar experiments using FPs would need a minimum of three colors (i.e., Histone H2B-CFP, a target kinetochore protein-EGFP, and a stable kinetochore marker-mCherry). However, overexpression of kinetochore proteins often causes mislocalization and mitotic defects^176–178^. This necessitates the generation of cell lines expressing endogenously-tagged proteins via CRISPR-Cas9 or similar techniques, which are time-consuming and impractical for large-scale studies of macro-molecular protein complexes, such as kinetochores. Additionally, compared to fluorescence dyes, FPs are significantly dimmer. For example, EGFP is at least 50% dimmer than Alexa 488 based on its extinction coefficient and quantum yield^179^. These FPs exhibit lower photostability and slower maturation. Furthermore, multiple fluorophore-conjugated secondary antibodies can bind to a single primary antibody, amplifying the signals far beyond what FPs can achieve. Together, these advantages of qIF enables highly sensitive and precise quantification of protein dynamics.

## Discussion

The kinetochore is a highly organized macromolecular protein complex composed of a diverse array of proteins, which are systematically and dynamically assembled throughout the cell cycle. However, a comprehensive understanding of kinetochore architecture during the cell cycle progression has remained elusive. In this study, we utilized qIF to precisely determine the dynamics of 36 different kinetochore proteins/substrates, covering key protein complexes and kinase substrates. Our findings provide a new insight into the dynamic architectural remodeling of kinetochores, shedding light on the assembly and disassembly of kinetochore components. Fig. 9 illustrates the proposed model of kinetochore architecture, integrating both our data and previous studies. In G1 phase, CENP-A, CENP-B and all CCAN proteins are consistently present at kinetochores. Notably, in the middle to late G1 phase, Mis12C is recruited to kinetochores via CENP-C. During the S phase, CCAN and Mis12C are rapidly assembled onto newly-synthesized centromeres, while sister kinetochores remain clustered within a sub-diffraction-limited distance until G2 phase. As Cyclin B1 levels rise during G2 phase^71^, Cdk1 becomes active, initiating the phosphorylation of key kinetochore substrates, such as CENP-C and CENP-T. Phosphorylation of CENP-C promotes its binding to CENP- A^36,40^, reshaping the CCAN architecture, including the dissociation of CENP-NL and CENP-HIKM complexes. Concurrently, phosphorylation of CENP-T facilitates its direct interaction with Ndc80C and Mis12C^58,180^. Additionally, the recruitment of Knl1C via Mis12C leads to the phosphorylation of MELT motifs on Knl1 by Mps1, triggering the assembly of the Bub1/Bub3 complex and their downstream proteins, including Sgo1, AurB, and Plk1. Although Mps1 begins accumulating at kinetochores during prophase^181^, its presence in the nucleus is detected as early as G2 phase^182^.

**Fig. 9.**
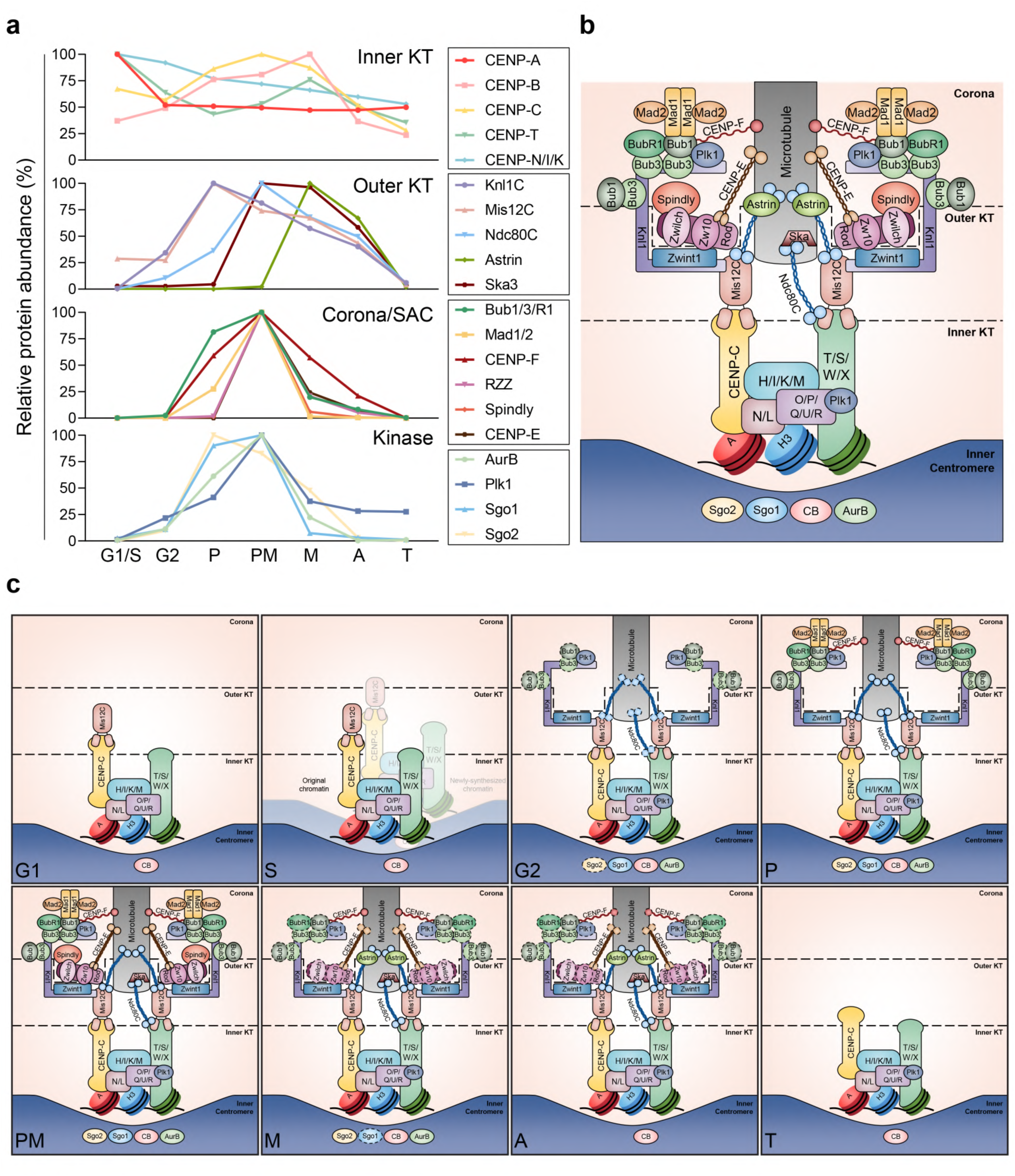
Summary of kinetochore landscape changes throughout the cell cycle. **a**, The relative abundance of individual kinetochore protein at kinetochores at each stage of the cell cycle. The data of each protein within the same complex was averaged. **b**, A model of the architecture of the human kinetochore. **c**, A model of the dynamic kinetochore landscape throughout the cell cycle.

During prophase, Cdk1, AurB, and Mps1 kinases reach their peak activities^181,183,184^, leading to the robust recruitment of Mis12C, Knl1C, and Bub1/Bub3 complex to kinetochores. This creates abundant binding platforms for SAC-related proteins. Simultaneously, Cdk1 and Mps1 cooperatively phosphorylate Bub1^96^, which facilitates Mad1 recruitment and the assembly of MCC^185,186^. The early assembly of MCC prior to NEBD suggests that pre-loading of SAC components to kinetochores may be critical to prevent premature chromosome segregation when microtubules begin to search and capture kinetochores immediately upon NEBD. Concurrently, AurB-mediated Bub1 phosphorylation triggers CENP-F recruitment to kinetochores^125^. After NEBD, proteins lacking NLS, such as the RZZ subunits and CENP-E, gain access to kinetochores. In early prometaphase, AurB reaches peak levels at kinetochores, and its phosphorylation of Zwint1 facilitates RZZ complex recruitment^113^, which in turn recruits Spindly and CENP- E^109,121^. In parallel, AurB intensifies CENP-F recruitment^125^. Additionally, Mps1 drives the oligomerization of the RZZ complex and Spindly^107,109,111^, forming a scaffold for the integration of Mad1, Mad2, CENP-E, and CENP-F, a process known as corona expansion. This phenomenon is particularly prominent in unattached kinetochores, as seen in nocodazole-treated mitotic cells^21^. As a result, most SAC-related and corona proteins reach their peak levels at this stage, enhancing the likelihood of kinetochore-microtubule interactions and promoting the transition from lateral to end-on microtubule attachment^11,118,187^. SAC strength is fine-tuned by the recruitment of BubR1, which peaks during prometaphase and enables the recruitment of the PP2A-B56 phosphatase to counterbalance Mps1 and Plk1 activities^86,89^.

During metaphase, chromosomes achieve biorientation, creating a physical separation between outer kinetochores and inner centromeres, where AurB is concentrated. This spatial arrangement facilitates the dephosphorylation of Knl1 by PP2A^83^, promoting the subsequent recruitment of PP1 to kinetochores. As a result, most of the MELT motifs become dephosphorylated, leading to the dissociation of Bub proteins^89^. Concurrently, end-on microtubule attachments strip off SAC and corona proteins, as evidenced by a dramatic reduction in RZZ complex, Spindly, Mad1/2, CENP- E, and CENP-F levels at kinetochores. However, a residual pool of Bub1, BubR1, RZZ, CENP-E, and CENP-F persists on metaphase kinetochores, suggesting the existence of two distinct protein populations: a tension-sensitive pool that dissociates upon chromosome biorientation, and a stable structural pool that likely contributes to the stabilization of kinetochore-microtubule attachments. Meanwhile, the reduction in AurB activity triggers Astrin-SKAP complex recruitment to kinetochores^168,170^. As cells progress into anaphase, Cdk1 and AurB activity declines further^62,64,184,188^, leading to a moderate reduction in all KMN network components and their downstream proteins, including the RZZ complex, CENP-E, and CENP-F. In telophase, all outer kinetochore, corona proteins, and mitotic kinases, except for Plk1, become nearly undetectable at kinetochores. The remaining Plk1 is responsible for CENP-A deposition onto centromeres during the early G1 phase^26^.

We found some interesting mismatches in the recruitment timing of kinetochore proteins, diverging from reported recruitment dependencies. These finding suggested the involvement of additional, yet unidentified regulatory mechanisms, potentially related to PTM, protein synthesis timing, or protein characteristics. Notable examples include the temporal gaps between Mad1 and its upstream adaptor Bub1, as well as Bub1 and BubR1. Despite the fact that both Bub1 and BubR1 bind to pMELT motifs, we observed a clear delay in BubR1 recruitment compared to Bub1. Although Mps1-mediated phosphorylation of Bub1 triggers its binding to Mad1^96^, a distinct time gap remains. Bub1 peaks at prophase and significantly reduces in prometaphase, while Mad1 levels increase (Fig. 4). Another example is the discrepancy between Zwint1 and RZZ complex. Zwint1 is recruited to kinetochores as early as prophase, but RZZ complex is not recruited until prometaphase. Given that Zw10 expression remains stable from G1/S to mitosis^189^, it is likely that RZZ complex is restricted from entering the nucleus until NEBD. In contrast, Spindly can enter the nucleus as early as G2 phase, but does not assemble at kinetochores until prometaphase when RZZ complex assembles at kinetochores. The precise mechanisms driving the time-gaps in kinetochore protein recruitment and dissociation remain unclear and warrant further investigation. In summary, our comprehensive qIF analysis elucidates the dynamic alteration of kinetochore landscape throughout the cell cycle. The insight gained from this study provides a valuable foundation for future research into these complex regulatory mechanisms.

## Acknowledgement

We would like to thank Yoshitaka Sekizawa and Yokogawa Electric Corporation for critical equipment and technical support. We would also like to thank Drs. Andrea Musacchio, Arshad Desai, Beth Weaver, Edward Salmon, Gary Golbsky, Iain Cheeseman, Jennifer DeLuca, Kinya Yoda, Mitsuhiro Yanagida, Tatsuo Fukagawa, Tomomi Kiyomitsu, Daniele Fachinetti, Song-Tao Liu, and Stephen Taylor for generously providing critical resources. Part of this work is supported by the Wisconsin Partnership Program Research Forward initiative from the University of Wisconsin-Madison Office of the Vice Chancellor for Research with funding from the Wisconsin Alumni Research Foundation, start-up funding from University of Wisconsin-Madison SMPH, UW Carbone Cancer Center, and McArdle Laboratory for Cancer Research, and NIH grant R35GM147525 and U54 AI170660 (to A.S.).

## Author contribution

YC.C. performed all experiments and analysis with the assistance of E.K., E.W., and W.R. A.S. conceptualized and supervised the entire project, contributing pivotal ideas and designing the experiments. YC.C. and A.S. prepared the manuscript draft. All authors reviewed and contributed to the manuscript’s refinement.

## Data availability

All data are available in the main text or the supplementary materials. Other data and original images used in this study are available from the corresponding author upon reasonable request.

## Methods

### Cell culture

Human RPE1 and HeLa cells were originally obtained from the American Type Culture Collection (ATCC, Manassas, VA, USA). CENP-C AID RPE1 cells^36^ were a kind gift from Dr. Daniele Fachinetti. Dsn1-AID RPE1 cells^175^ were a kind gift from Dr. Tatsuo Fukagawa. All above cell lines were cultured in DMEM (Gibco, 11965092) or DMEM/F12 (Gibco, 11320033) supplemented with 1% penicillin-streptomycin and 10% fetal bovine serum (Seradigm, 1500-500) under 5% CO_2_ at 37°C in an incubator. For inhibitor treatments, cells were treated with 10 μΜ MG-132 (MCE, HY-13259), 3 μΜ nocodazole (Thermo Fisher Scientific, AC358240100), 2.5 or 5 μΜ STLC (Sigma-Aldrich, 164739), 10 μΜ ZM447439 (MCE, HY-10128), or 1 μΜ Palbociclib (MCE, HY-50767). 500 μΜ of Auxin (Sigma-Aldrich, 12886) was used for CENP-C-AID RPE1 cells and Dsn1-AID RPE1 cells. For the cold-stability assay, cells were incubated in cold media at 4°C for 10 min to depolymerize unstable spindle fibers before fixation.

### Immunofluorescence

Asynchronous RPE1 cells were seeded on #1.5 thickness coverslips at least a day prior to fixation. Cells were fixed by 4% PFA in 250 mM HEPES buffer (pH 7.5) at 37°C or cold methanol at −20°C for 15 min. For Bub3 and Ska3 stanning, pre-extraction was performed by incubation of cells with 1% Triton X-100 in PHEM buffer at 37°C for 1 min, followed by fixation with 1% Triton X-100 in 4% PFA/HEPES at 37°C for 15 min. After fixation, cells were washed with PBS 3 times at room temperature (RT). Only cells fixed with PFA were then permeabilized by 0.5% Nonidet P-40 (Santa Cruz, sc-29102) in PBS at RT for 15 min. After permeabilization, cells were washed with PBS. Blocking was performed by incubation of cells in 0.1% Bovine Serum Albumin (Sigma-Aldrich, A2153). Cells fixed with PFA were incubated in primary antibody solution in a humidified chamber at 37°C followed by incubation in secondary antibody solution in a humidified chamber at 37°C. Cells fixed with methanol were incubated in primary antibody solution in a humidified chamber at RT, followed by incubation with secondary antibody solution in a humidified chamber at RT. Primary and secondary antibody-related information are listed in Supplementary Table 1-3. After secondary antibody staining, cells were washed with PBS. Cells were incubated in 200 ng/ml DAPI (Sigma-Aldrich, 13190309) in PBS at RT for 15 min. The stained coverslips were mounted with homemade mounting media (20 mM Tris (pH 9.0), 90% glycerol, 0.2% n-propyl gallate).

### Imaging

Images were acquired using Nikon Ti2 inverted microscope equipped with Yokogawa CSU-W1 spinning disc confocal and Hamamatsu Quest qCMOS camera or Nikon Ti2 inverted microscope equipped with Yokogawa CSU-SoRa W1 spinning disc confocal and Hamamatsu Fusion camera. Both microscopes were equipped with a high-power laser unit (100 mW for 405, 488, 561, and 640 nm wavelength) for excitation. Z-stack images were acquired at a step of 0.2 μm controlled by Nikon NIS Elements (version 5.21). Plan Apo 100x oil objective (NA = 1.45) was used for qIF assay, and Plan Apo λ 60x oil objective (NA = 1.40) was used for other experiments. Images of all cell cycle stages for one biological replicate were acquired from the same single coverslip on the same day using the same imaging settings. For all qIF assays, the representative images of all cell cycle stages using the same set of antibodies were adjusted to have the same brightness and contrast for the target protein as a fair comparison.

### Image analysis

The local background corrected signal intensity measurement for a single kinetochore was described in the previous research^20^ and a schematic representation was shown in Extended Data Fig. 1d,e. Signal quantification was performed using MetaMorph (version 7.10). Briefly, two bounding boxes with different sizes were placed on the target kinetochore at the best focus z plane. The area between two boxes was used to determine the local mean background intensity. The true signal intensity of the target kinetochore was calculated by subtracting the local background intensity for each kinetochore. Since the diameter of a kinetochore is about 250 nm, only the best focus single z plane (the z plane with highest maximum intensity) was used to determine the signal intensity of individual kinetochores. To quantify the percentage of positive cells with kinetochore signals, cells were co-stained with antibodies for target proteins, including Dsn1, Pmf1, CENP-T, CENP-N, and CENP-I, and a kinetochore marker, either CENP-C or ACA. Cells lacking detectable levels of target protein signals at kinetochores were considered negative cells.

### Statistics and biological replicates

All quantification plots were made using GraphPad Prism (version 9.5). All statistics in this study were performed with unpaired Student’s t-test or Tukey’s multiple comparisons test using GraphPad Prism. All experiments had 2-3 independent biological replicates performed. Sample size and the number of biological replicates were included in each figure. Each data point represents a single kinetochore, except for some parts of the dataset of CENP-B, Sgo1, Sgo2, and AurB as well as the entire measurements for pH2- T120 and pH3-T3. For CENP-B, Sgo1, Sgo2, and AurB, each data point from G2 phase to metaphase reflected the combined signal intensity of paired sister kinetochores. In the case of pH2A-T120 and pH3-T3, each data point represents the signal intensity within a single nucleus. Kinetochore signal intensities at each cell cycle stage were normalized to the stage exhibiting the highest mean signal intensity across the cell cycle stages. All error bars represent standard deviation relative to the mean, and independent replicates are distinguished by different colors.

